# Genomic health is dependent on long-term population demographic history

**DOI:** 10.1101/2022.08.16.503900

**Authors:** Eric Wootton, Claude Robert, Joëlle Taillon, Steeve Côté, Aaron B.A. Shafer

## Abstract

Current genetic methods of population assessment in conservation biology have been challenged by genome-scale analyses due to their quantitatively novel insights. These analyses include assessments of runs-of-homozygosity (ROH), genomic evolutionary rate profiling (GERP), and mutational load. Here, we aim to elucidate the relationships between these measures using three divergent ungulates: the white-tailed deer, caribou, and mountain goat. The white-tailed deer is currently expanding, while caribou are in the midst of a significant decline. Mountain goats remain stable, having suffered a large historical bottleneck. We assessed genome-wide signatures of inbreeding using the inbreeding coefficient *F* and %ROH (*F_ROH_*) and identified evolutionarily constrained regions with GERP. Mutational load was estimated by identifying mutations in highly constrained elements (CEs) and sorting intolerant from tolerant (SIFT) mutations. Our results show that *F* and *F_ROH_* are higher in mountain goats than in caribou and white-tailed deer. Given the extended bottleneck and low *N_e_* of the mountain goat, this supports the idea that the genome-wide effects of demographic change take time to accrue. Similarly, we found that mountain goats possess more highly constrained CEs and the lowest dN/dS values, both of which are indicative of greater purifying selection; this is also reflected by fewer mutations in CEs and deleterious mutations identified by SIFT. In contrast, white-tailed deer presented the highest mutational load with both metrics, in addition to dN/dS, while caribou were intermediate. Our results demonstrate that extended bottlenecks may lead to reduced diversity and increased *F_ROH_* in ungulates, but not necessarily the accumulation of deleterious alleles, likely due to the purging of deleterious alleles in small populations.

## Introduction

Organizations like the International Union for Conservation of Nature (IUCN) rely primarily on population sizes and trends when assigning wildlife conservation statuses (IUCN Standards and Petitions Committee, 2019). The predictive relationship between population size (*N_C_*) and genetic diversity is challenging, as *N_C_*, for example, tends to be substantially more variable than diversity metrics like nucleotide diversity (*π*), referred to as Lewontin’s paradox (Ellegren & Galtier, 2016; Lewontin, 1974), and therefore rarely accounts for recent demographic history and the genetic health of populations. While genetic diversity can correlate with IUCN status (Peart et al., 2020; Petit-Marty et al., 2021), the relationship (i.e., direction) is varied and predictive association appears low (Schmidt et al., 2022). The unprecedented loss of global biodiversity (Pascual et al., 2021) and questionable genetic diversity associations (Schmidt et al., 2022), would strongly suggest that traditional conservation genetic markers and indices must be replaced with more universally informative metrics, if they in fact exist.

Both novel genome metrics and functional versus presumed neutral variation have garnered recent attention in the conservation genetics community. The use neutral genetic diversity in conservation biology has a long history (e.g. DeWoody et al., 2021; Kohn et al., 2006), with for example specific targets of management focused on retaining 90% of initial [neutral] genetic diversity (Ballou & Foose, 2010). With the advent of whole-genome sequencing, the notion of targeted gene flow (Kelly & Phillips, 2019) and gene-editing (Phelps et al., 2020) for the purposes of conservation emerged. Teixera & Huber (2021) further suggested the focus be shifted to functional variation and gene-environment interactions, in part due to the poor associations to IUCN status noted above. Gene-targeted approaches, however, come with a suite of potential undesirable outcomes (Kardos & Shafer 2018), and genome-wide diversity still appear to be the most indicative of population viability (Kardos et al. 2021). Likewise, Peart et al. (2020) showed that genome-wide estimates of Tajima’s *D*, a composite statistic derived from Watterson’s *θ* and *π* (Tajima, 1989), appears predictive of conservation status. Importantly, the uncertainty around which genetic marker to use reflects both a shifting understanding and nuanced patterns of genomic diversity, and further reinforces the need for identifying less ambiguous conservation-informative genetic metrics.

The general correlation between genome-wide diversity and microsatellite heterozygosity at the population-level suggests that re-sequencing data reflect smaller targeted panels of loci (Väli et al., 2008). This observation is ultimately what formed the premise of Shafer et al. (2015) that argued for the use of quantitatively novel genomic metrics for conservation. Runs-of-homozygosity (ROH), genomic evolutionary rate profiling (GERP), and mutational load are metrics dependent on genome-scale data and have only recently been applied to species and populations of concern (Grossen et al., 2020; van der Valk et al., 2019; von Seth et al., 2021). ROHs are long stretches of homozygosity that arise as a result of inbreeding (Ceballos et al., 2018), and reflect the true inbreeding coefficient and can identify genes underlying inbreeding depression (Kardos et al., 2016). GERP profiling can be used to identify evolutionarily constrained genomic elements, defined as regions with fewer substitutions than expected under neutral evolution (Davydov et al., 2010), which are often the subject of selective pressures (Cooper et al., 2005; Davydov et al., 2010). Any mutations that appear at highly conserved sites are generally assumed to be deleterious (Cooper et al., 2005; Davydov et al., 2010). Likewise, tolerant and intolerant mutations – also referred to as SIFT scores (Vaser et al., 2015) – that are considered of human clinical interest (P. Kumar et al., 2009) – have now been applied to natural populations (Robinson et al., 2016). ROH, GERP and SIFT scores collectively allow for inferring the mutational load and genomic health of a population or species.

While quantitatively useful, the relationships between these metrics and conservation status and diversity metrics are generally not well defined and at-times counter to expectation. van der Valk et al. (2019) showed a weak relationship between conservation status and mutational load across mammalian taxa; the highest mutational loads were in fact seen in the most abundant and diverse species. Similarly, species with the highest inbreeding (ROH) also had the lowest mutational load and GERP scores (van der Valk et al., 2019). This pattern could be due the purging of deleterious alleles in small populations, which is consistent with simulations by Kyriazis et al. (2021). van der Valk et al. (2019) argued that this process has implications for genetic rescue and suggested that individuals be selected for translocation based on low mutational load and not diversity. Work on the Sumatran rhinoceros (*Dicerorhinus sumatrensis*) added another layer of nuance, with populations showing low to moderate levels of inbreeding (ROH), but a relatively high mutational load (von Seth et al., 2021). In contrast, the Alpine ibex (*Capra ibex*) that has experienced successive demographic bottlenecks leading to reduced diversity and increased ROH, has not accumulated highly deleterious alleles (Grossen et al., 2020). Thus, there seems to be a link between historical demography, ROH, and mutational load. Understanding such processes has important implications for both predicting long-term viability, but also selecting animals for genetic rescue.

Here, we analysed signatures of genomic health across populations of three divergent ungulates to elucidate these relationships. The caribou (*Rangifer tarandus*) is currently experiencing significant population declines (*N_C_*) prompting international concern (COSEWIC, 2015). In Quebec, five of the major populations have seen drastic declines from their highest estimates ranging between 10% and 99%, depending on the herd (COSEWIC, 2015, 2017). Historical estimates of *N_e_* suggest fluctuations tracking glacial cycles, with more recent estimates in the thousands (Taylor et al. 2021). The white-tailed deer (*Odocoileus virginianus*), in contrast, is expanding northward and has accordingly high genomic diversity (Kessler et al., 2022). While white-tailed deer currently number in the 10s of millions (Kauffman et al., 2018) with historic *N_e_* estimates in the tens of thousands (Lamb et al. 2021), deer were extirpated across much their range due to colonial harvest and forestry practices (Kauffman et al., 2018); meaning this demographic expansion is very recent. The mountain goat (*Oreamnos americanus*) underwent a massive bottleneck during the last glacial maximum with *N_e_* estimates in the hundreds (Martchenko et al., 2020; Martchenko & Shafer 2022); mountain goats have since remained relatively stable with a range-wide population of ~100,000 individuals (Festa-Bianchet & Côté, 2008). These three species, contrasted in their ecology but also conservation status, historical and demographic trajectories, present useful comparisons to explore patterns of novel genomic metrics as they pertain to conservation status. In line with recent empirical work, we predicted that populations with low diversity and high inbreeding do not necessarily have higher mutational load and that genome-wide effects of demographic change like ROH take time to accrue. Similarly, we expected to see greater purging of deleterious alleles in populations with long-term small *N_e_*. Collectively, this means that poor genomic metrics of health would not correlate with current negative population trends.

## Methods

Caribou samples were obtained from 67 individuals across Quebec and Labrador (Dedato et al. 2022); 53 white-tailed deer were sampled across their North American range (Kessler et al. 2022). Additionally, 35 mountain goats were sampled across their native range from Alaska to as far south as Idaho (Martchenko et al. 2022; Figure 1). All samples were sequenced with Illumina HiSeqX technology to approximately 5X coverage. Non-target sequences were trimmed from raw FASTQ files with Trimmomatic v.0.36 (Bolger et al., 2014), and reads were mapped to their reference genome using BWA-MEM v.0.7.17 and SAMtools v.1.15 (Li et al., 2009; Li & Durbin, 2009). Duplicate mapped reads were marked and removed using Picard v.2.23.0 (Broad Institute, 2019). Uniquely mapped reads were subset by Sambamba v.0.8.2 (Streit et al., 2012), and indels were realigned with GATK v.3.8 (Van der Auwera et al., 2013). We estimated genotype likelihoods using ANGSD v.0.934 and the GATK model: -doVcf 1 -doGeno -4 -doPost 1 -gl 2 -SNP_pval 1e-6 -minMapQ 20 -minQ 20 -doCounts 1 -skipTriallelic 1 -doGlf 2 (Korneliussen et al., 2014). Hard calls required for analyses were performed by Beagle v.5.1 (Browning et al., 2018) with the following parameters: impute=false overlap=3000 window= 100000 gprobs=TRUE parameters. Posterior genotype probabilities were estimated using allele frequencies, and only genotypes with a posterior probability above 0.95 were called. Tajima’s *D* and nucleotide diversity (π) were calculated using VCFtools v.0.1.13 with windows of 10 Kb and using all positions, regardless of allele frequency (Danecek et al., 2011).

**Figure 1:**
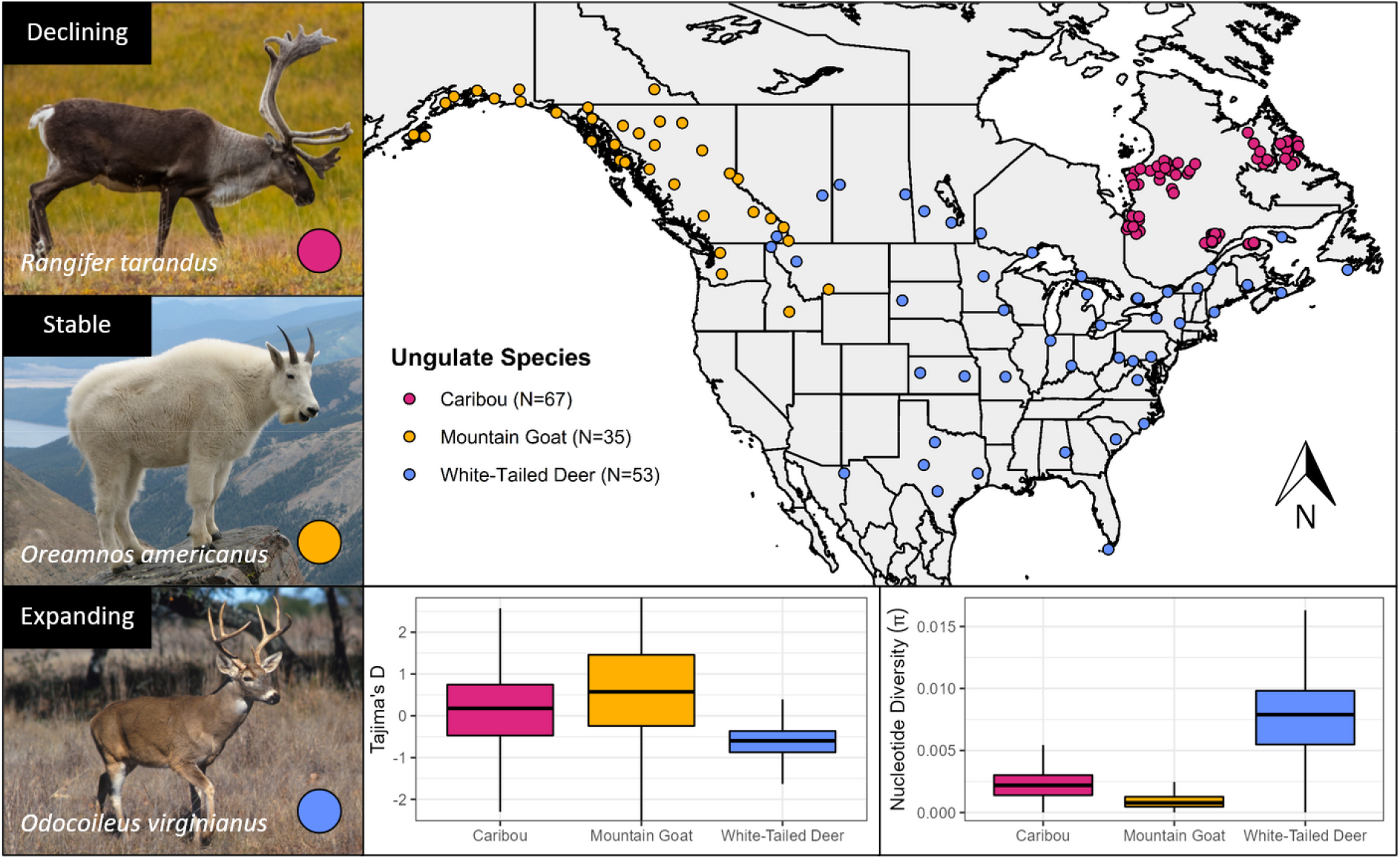
Sample locations and images of each species. (Red) *Rangifer tarandus* (N=67), (Yellow) *Oreamnos americanus* (N=35), (Blue) *Odocoileus virginianus* (N=53). Current demographic trends are outlined in the top left corner of each image. Boxplots showing Tajima’s D and nucleotide diversity (π) per species are also pictured.

### Inbreeding and Homozygosity

A subset of SNPs in approximate linkage equilibrium was obtained using PLINK v.1.9 with a window size of 50 SNPs, a window shift size of 10 SNPs, and a pairwise (R^2^) linkage disequilibrium threshold of 0.1 (Chang et al., 2015; Purcell et al., 2007). We then identified runs of homozygosity (ROH) on scaffolds greater than 10 Mb in length and with a minor allele frequency cut-off of 0.05 (Foote et al., 2021). Sliding windows of 50 SNPs were used, and each ROH was defined as a sequence at least 100 Kb long with at least 50 homozygous SNPs (Foote et al., 2021). Each ROH required a minimum density of 1 SNP per 50 Kb and no more than 10 missing or 3 heterozygous calls in a window. Any SNPs more than 1 Mb apart within a segment were used to split that segment in two. Once identified, each ROH was classified by size, and the percent ROH (*F*_ROH_) for each individual was calculated. PLINK parameters were taken from Foote et al. (2021) with output validated using the program ROHan which identifies ROHs using genomic windows of fixed size (Renaud et al., 2019). ROHan was run on the same data with the same ROH size definition of 100 Kb and a transition to transversion ratio (Ts/Tv) of 2.25. This ratio was calculated using BCFtools v.1.14 and the genotypes provided by ANGSD (Li et al., 2009). The inbreeding coefficient (*F*) for each individual was also estimated using PLINK and the –het flag.

### Evolutionary Constraint and Mutational Load

Twenty ungulate reference genomes from the NCBI GenBank genome database (Table S1) were selected to represent most extant families with genomes available in both the orders of Perissodactyla (odd-toed ungulates) and Artiodactyla (even-toed ungulates). Constrained regions for each individual of our three focal species were identified using the Genomic Evolutionary Rate Profiling (GERP++); this approach requires a phylogenetic tree which we constructed for all 23 species using TimeTree (Figure S1) (Kumar et al., 2017). The raw paired-end reads of all study individuals of the were then aligned to each of the 20 reference genomes in 50 bp segments using BWA-MEM to produce 20 alignment files per individual (3100 in total). All duplicate alignments were marked and removed with Picard. Then, the alignments of each study individual were concatenated into one multiple alignment file (MAF) in pileup format using HTSbox v.1.0, resulting in 155 total MAFs (one per individual). This method of alignment uses each study individual as a focal reference and was first reported by von Seth et al. (2021). To eliminate bias, each focal species was omitted from the MAF before running GERP++. The gerpcol function included in GERP++ was run on each MAF to generate rejected substitution (RS) scores for each alignment column. RS scores are used as a metric for constraint intensity and are calculated by subtracting the estimated evolutionary rate at each position from the neutral rate (Cooper et al., 2005; Davydov et al., 2010). Both rates were estimated by gerpcol using the aforementioned phylogenetic tree (Figure S1). RS scores quantify the number of differences between the expected and observed number of SNPs at a site, and a positive score is indicative of constraint (Cooper et al., 2005; Davydov et al., 2010). An RS score greater than 2 represents a highly constrained region whereas a score lower than 1 is putatively neutral and discarded (Cooper et al., 2005; Davydov et al., 2010). gerpcol was run with the previously calculated Ts/Tv ratio of 2.25. We also used GERP++ to identify constrained elements with the gerpelem function.

To estimate mutational load, we determined the number of SNPs called by Beagle within these constrained elements using BEDtools v.2.30.0 (Quinlan & Hall, 2010). Here the ancestral allele at each site is considered the variant present in the phylogenetically closest outgroup (Fig. S1), allowing derived alleles to be inferred. These derived SNPs are putatively deleterious due to the common functional importance of constrained regions (Cooper et al., 2005; Davydov et al., 2010). For each species, the functional consequence of each SNP called by Beagle was determined using Ensembl’s Variant Effect Predictor (VEP) with default parameters, corresponding species reference genomes, and GFF format annotation (McLaren et al., 2016; Table S1). The deleteriousness of each SNP across the genome was then estimated using functional tolerance scores provided by SIFT4G with default parameters (Vaser et al., 2015). Using the reference genomes and annotation for each study species alongside UniRef90 (Suzek et al., 2015), protein databases were generated by SIFT for each species. SIFT calculates normalised probabilities using these databases where substitutions with scores lower than 0.05 are predicted to be deleterious to protein function (Vaser et al., 2015).

The pN/pS ratios for each individual were calculated as the ratio of nonsynonymous to synonymous mutations identified by SIFT. Likewise, following Jeffares et al. (2015), the CODEML application from PAML v.4.9 was used to estimate dN/dS for each individual (Yang, 2007). We filtered each of the MAFs used in the GERP analysis to include only coding regions using the extractcds program from the EMBOSS suite of software (Rice et al., 2000). The positive selection model (M2a) was used in CODEML to then analyse each MAF with the phylogenetic tree from TimeTree.

Caribou did not have an available annotation at the time of analysis which is required for these analyses; thus, we used a mixture of *de novo* and homology-based predictions. We aligned protein sequences from *Bos taurus, Equus caballus, Homo sapiens*, and *Ovis aries* to the caribou genome using blastx v.2.7.1 (Altschul et al., 1990). These sequences were all obtained from Ensembl release 89 (Hunt et al., 2018). We also aligned transcripts from the NCBI mammalian RefSeq 10ranscript database v.1.0 using BLAT v.1.0 to help identify exons. Caribou RNAseq data (SRX12486288) from the NCBI Sequence Read Archive were aligned to the genome with HISAT v.2.10 for *de novo* prediction (Kim et al., 2015). Using these data, genes were predicted by Augustus v.3.1.1 as well as GeneMarkES v.4.69 (Lomsadze et al., 2005; Stanke et al., 2006). We generated a set of consensus genes using EvidenceModeler v. 1.1.1 with the following weighting scheme (Haas et al., 2008): transcript alignment via BLAT (10x weight); protein alignment via blastx (5x weight); gene prediction via Augustus (1x weight); gene prediction via GeneMarkES (1x weight). PASA v.2.3.3 was used last to refine gene identifications (Haas et al., 2008).

For all individual metrics above we quantified the correlation to sample location (latitude, longitude). Because white-tailed deer were extirpated across much of their range, we used translocation data from McDonald & Miller, (2004) to determine if these metrics were correlated to restocking efforts in the United States. We examined correlations with stock start and end dates, estimated population lows, number of individuals stocked, and the number of states involved in stocking.

## Results

### Diversity and Inbreeding

Nucleotide diversity (*π*) was found to be highest in deer (mean = 0.00771) and lowest in mountain goats (mean = 0.00101) (Figure 1). Tajima’s *D* was highest in mountain goats (mean = 0.625) and lowest in white-tailed deer (mean = −0.645) (Figure 1). For caribou, Tajima’s D was positive (mean = 0.124), but lower than in mountain goats (t-test p-value < 2.2×10^-16^). Mountain goats were found to have significantly higher *F_ROH_* (mean = 0.234) than both caribou (0.00184) and white-tailed deer (mean = 0.00380) (Figure 2a). This pattern is reflected in the relative values of *F* where mountain goats had the highest value (mean = −0.018) in comparison to caribou (mean = −0.44) and deer (mean = −0.27) (Figure 2b). For the mountain goat, most ROHs were found to be in the 500-1000 Kb range whereas caribou and deer possessed ROHs almost exclusively below 300 Kb (Figure 3). Despite a much higher number of ROHs overall, mountain goats possessed significantly fewer in the 100-300 Kb range than caribou (Figure 3). Both measures of inbreeding were found to be correlated in mountain goats (Pearson’s *r* = 0.918, *p* < 0.05), deer (Pearson’s *r* = 0.213, *p* < 0.05), and caribou (Pearson’s *r* = 0.199, *p* < 0.05) (Figure S2).

**Figure 2:**
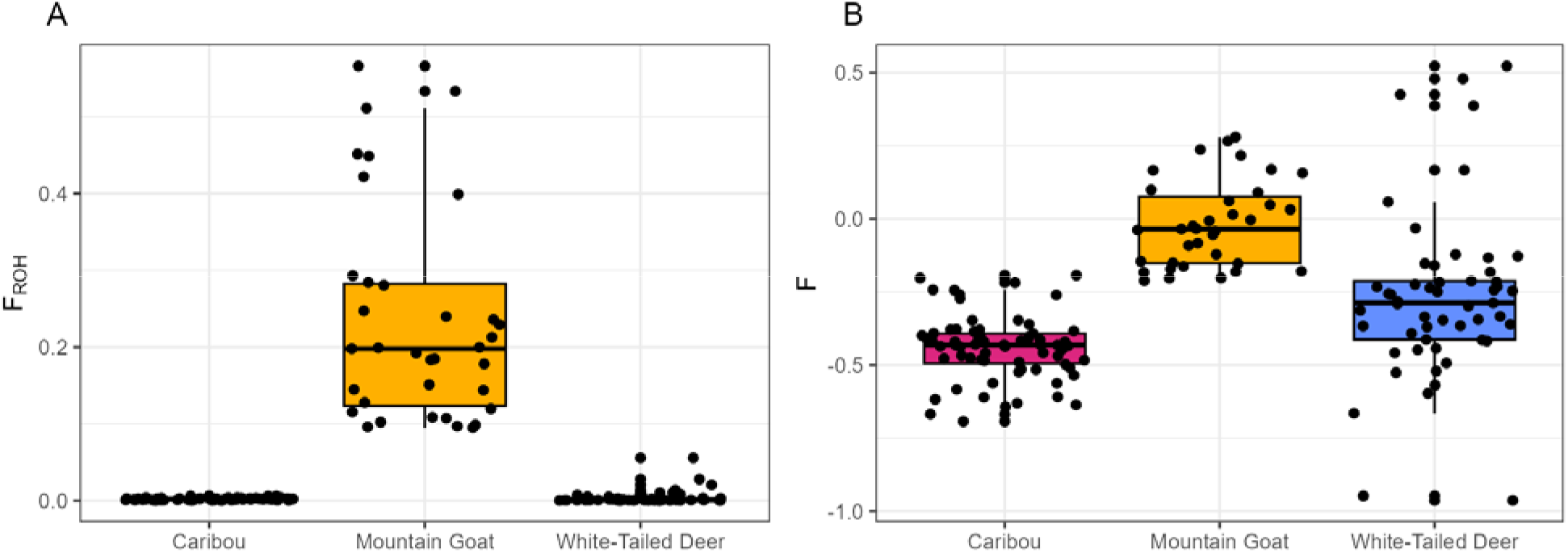
(A) Boxplot showing *F_ROH_* as calculated by PLINK for all three species. Only ROHs greater than 100 Kb are included in each estimate. (B) Boxplot showing the method-of-moments inbreeding coefficient (*F*) for each species as calculated by PLINK.

**Figure 3:**
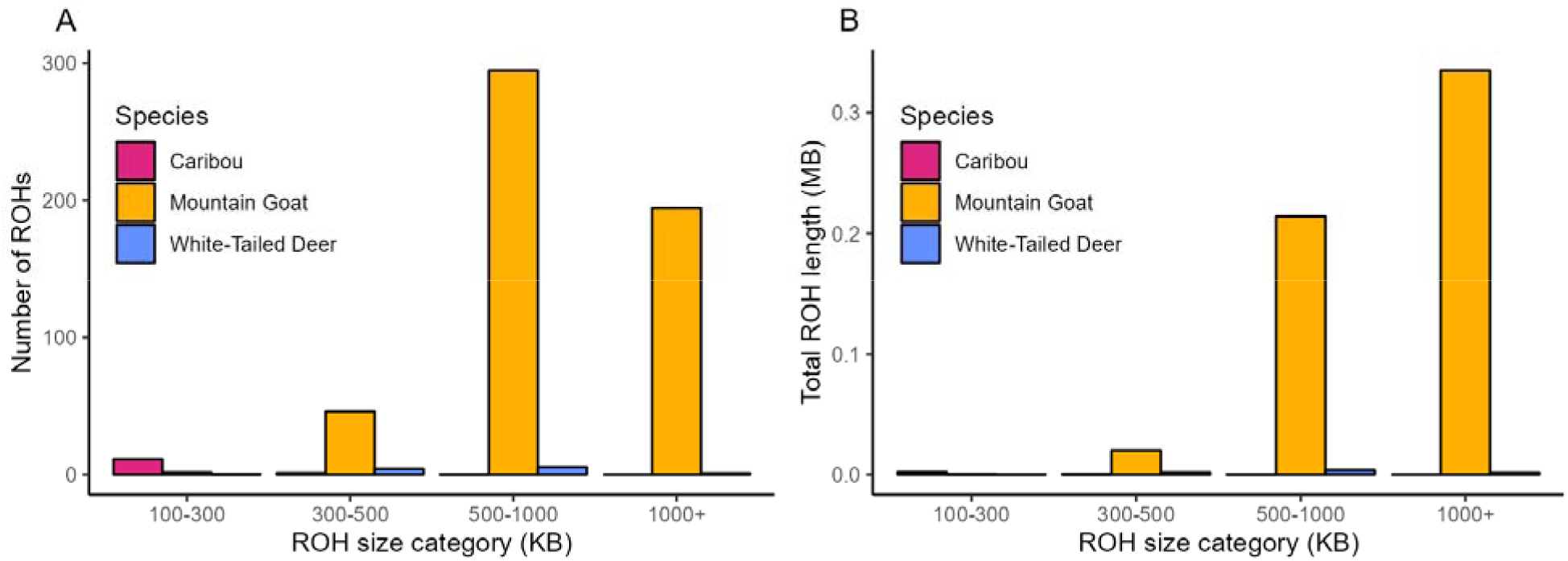
(A) Bar plot showing the total number of ROHs in each species stratified by ROH size category in Kb. (B) Bar plot showing the total length (Mb) of all ROHs in each size category (Kb).

### Evolutionary Constraint and Mutational Load

Mountain goats had the highest number of constrained elements (Ces) (mean = 2926) while caribou have the fewest (mean = 935; Figure S3). Highly constrained elements followed a nearly identical distribution between species with all three having approximately 50% as many Ces with RS > 2 (Figure S3). In addition to having the most Ces, mountain goats showed slightly longer Ces on average (Figure S4). The three species showed little variability in average CE RS-score (Figure S4, S5). Despite having significantly fewer Ces, white-tailed deer had a larger number of SNPs in Ces (mean = 411) followed by mountain goats (mean = 212) and caribou (mean = 74) (Table 1). The relative differences between species remained the same for highly constrained Ces (Table 1). We observed that a much higher fraction of constrained positions was deleterious in deer (mean = 0.625%) than in caribou (mean = 0.326%) and mountain goats (mean = 0.121%; Figure 4a). The pN/pS ratio for each species was also highest in deer (mean = 0.446) and lowest in caribou (mean = 0.175) and mountain goats (mean = 0.182) (Figure S6); dN/dS for each species followed a similar distribution with deer having much larger ratios (mean = 0.645) than caribou (mean = 0.166) and mountain goats (mean = 0.103) (Figure 4b).

**Figure 4:**
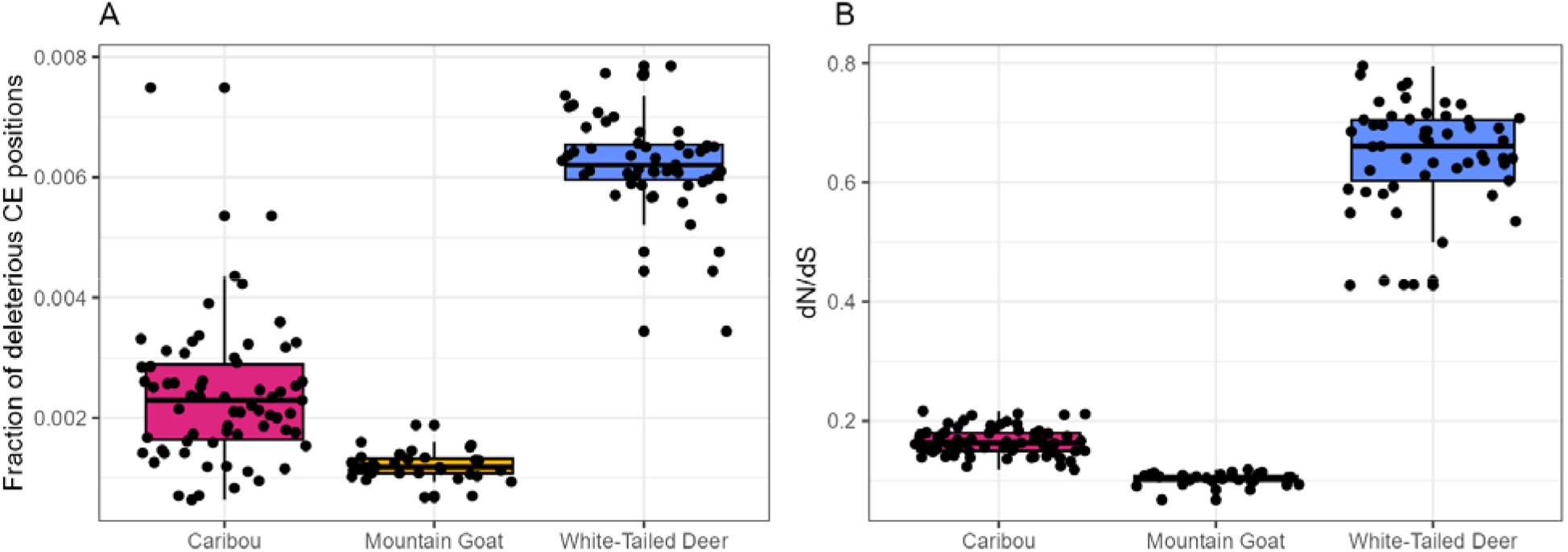
(A) Boxplot showing the fraction of constrained positions with SNPs by species. Such SNPs are putatively deleterious, making this the fraction of positions that are deleterious in constrained elements (CEs). (B) Boxplot showing dN/dS calculated for each individual by CODEML.

**Table 1:** The number of scaffolds > 10 Mb, mean scaffold length > 10 Mb, number of genes, and average number of deleterious mutations per species identified by GERP and SIFT (total and homozygous). The number of genes was taken from the annotation of each species.

| Species | Caribou | Mountain Goat | White-Tailed Deer |
| --- | --- | --- | --- |
| Scaffolds > 10 Mb (n) | 68 | 49 | 86 |
| Mean scaffold length > 10 Mb | 19,723,800 | 49,667,287 | 21,282,945 |
| Genes (n) | 33,242 | 22,292 | 20,750 |
| SNPs in CEs (GERP) | 74 | 212 | 411 |
| SNPs in CEs with RS > 2 (GERP) | 32 | 74 | 187 |
| Deleterious mutations (SIFT) | 4,943 | 3,190 | 6,834 |
| Homozygous deleterious mutations (SIFT) | 2,509 | 1,701 | 3,570 |

We found 42.1% of coding sequence consequences identified by VEP to be missense variants in white-tailed deer, 49.3% in caribou, and 57.4% in mountain goats (Figure 5a). Despite this distribution, white-tailed deer had the highest number of intolerant mutations (mean = 6852) identified by SIFT followed by caribou (mean = 4944) and mountain goats (mean = 3190) (Table 1). Similarly, we observed white-tailed deer to have the highest ratio of deleterious mutations to genes (mean = 0.330) while caribou (mean = 0.149) and mountain goats (mean = 0.143) had the lowest (Figure 5b). None of these metrics were correlated to the number of genes in the annotation (Table 1).

**Figure 5:**
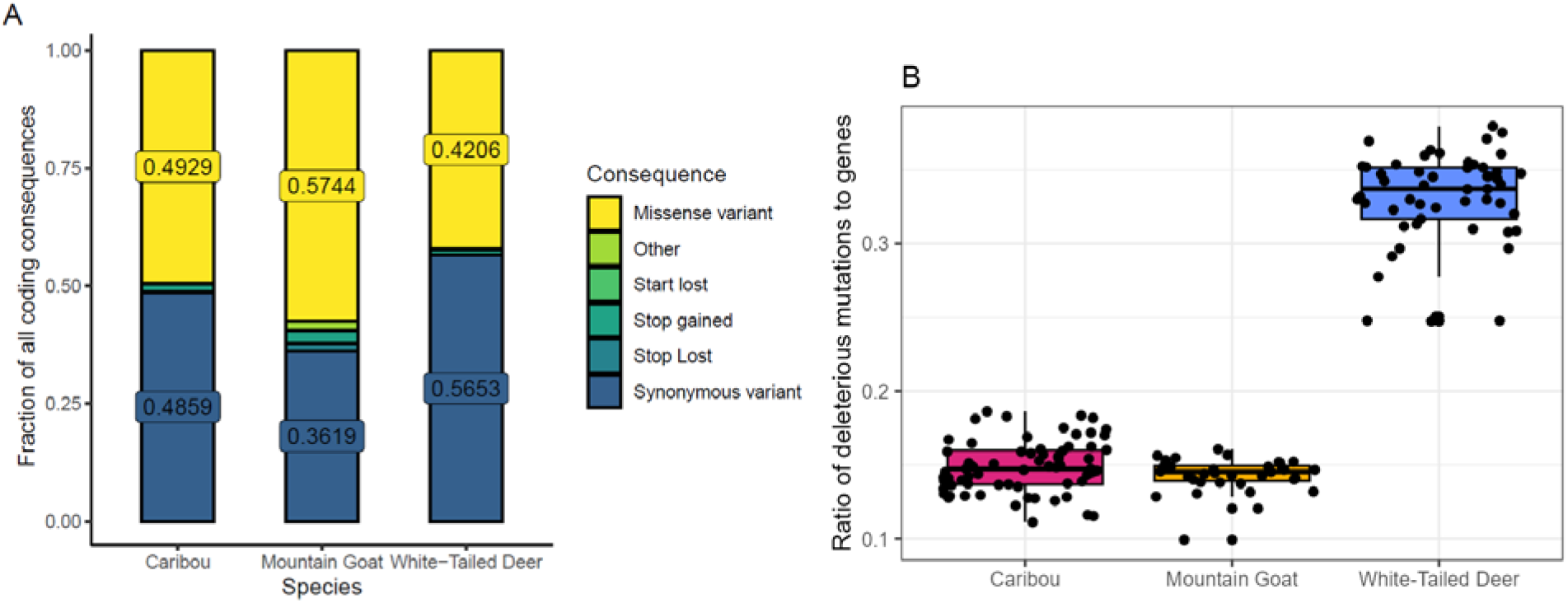
(A) Bar plot showing the fractions of each coding consequence identified by VEP in each species. (B) Boxplot showing the ratio of deleterious mutations identified by SIFT to the number of genes in each species.

Lastly, we found *F* and latitude to be moderately correlated in caribou (Pearson’s *r* = −0.466, *p* < 0.05; Figure S7) and deer (Pearson’s *r* = −0.513, *p* < 0.05; Figure S7), but not in mountain goats (Pearson’s *r* = 0.003, *p* < 0.05). There were no significant correlations between the individual genome health metrics and restocking data in white-tailed deer (Pearson’s *r* < 0.4, *p* > 0.05; Table S2).

## Discussion

The interplay between nucleotide diversity (π), inbreeding, mutational load, and census size (*N_C_*), is nuanced and full of contrasting examples. Here, we demonstrate that mutational load may be lower than expected in populations with relatively high inbreeding and low diversity. Our observed patterns suggest that the purging of deleterious alleles in these cases is stronger in small populations (Huber et al., 2020), and therefore would reduce risks from inbreeding depression. This supports our hypotheses that low genetic diversity and evidence of inbreeding do not on their own necessitate special conservation status, and that purging is likely occurring in populations with low *N_e_*. We also predicted that genomic signatures of demographic events take time to accrue, which is supported by evidence of *F_ROH_* and CEs requiring extended bottlenecks to surface and increase. Caribou have seen recent decreases in *N_C_* yet have no detectable ROH; in contrast, mountain goats show elevated ROH, likely due to the ancestral nature of their bottleneck (Martchenko et al., 2020). The varied relationships between different measures of genomic health and the lack of correlation to the population trends that are typically used for conservation status assignment highlights the value of this comparative framework (see also Huber et al., 2020). This information is especially valuable for small populations of conservation concern, where mutational load combined with ROH is probably the most informative way forward, in addition to metrics like Tajima’s *D* (Peart et al., 2020).

### Inbreeding Effects

Genomic consequences stemming from demographic change take time to accumulate (e.g. Grossen et al., 2020; von Seth et al., 2021). Historical demographic trends inferred from SMC approaches suggest both caribou and mountain goats responded negatively to the last glacial maximum (Taylor et al. 2021; Martchenko et al. 2020). More recent estimates (<10 KYA) show mountain goats with a considerably smaller *N_e_* (≪1000) compared to caribou (Martchenko & Shafer 2022, Taylor et al. 2020). In contrast, demographic trends in white-tailed deer present as a much more subtle response to glacial cycles, with a noticeably higher recent *N_e_* in the tens of thousands (Haworth et al., 2021; Lamb et al., 2021). Variation in temporal *N_e_* almost certainly stems from current white-tailed deer range being below the ice sheets, suggesting they were less impacted, which was not the case for mountain goats and caribou. Importantly, these SMC-derived patterns of *N_e_* also mirror our estimates of *π* (Figure 1); this is consistent with long-term estimates of *N_e_*, derived from *π* = *4N_e_μ*, matching the averaged temporal PSMC estimates of *N_e_* (Peart et al. 2020).

Caribou populations are currently in the midst of a demographic bottleneck resulting in a low contemporary effective and census population size (*N_e_*; Dedato et al., 2022). In contrast, mountain goats underwent a large historical bottleneck during the last glacial maximum, and while they have not recovered, they have remained relatively stable, resulting in a low *N_e_* (Martchenko & Shafer, 2022). Low diversity stemming from bottleneck is supported by positive values of Tajima’s *D* in these two species, which indicate a lack of rare alleles and population contraction (Tajima, 1989). The lower *π* in mountain goats is clearly due to their extended bottleneck that would have exacerbated any effects of inbreeding, but also their historically lower census sizes. These effects are reflected by *F_ROH_*, where mountain goats were found to have 23.4% of the genome in ROH whereas caribou have < 1%. These differences are also unlikely to be the result of genome assembly quality, as the mean number and length of analysed scaffolds were comparable (Table 1).

Compared to caribou and white-tailed deer, mountain goats have developed significantly longer and more numerous ROHs, which further outline the effects of a prolonged bottleneck. Caribou simply have not been subject to a low *N_C_* for long enough to approach mountain goats or even differentiate themselves from deer in their levels of inbreeding, with for example explicit demographic models detecting historical, not recent bottlenecks (Dedato et al. 2022). Of note, *F_ROH_* was more correlated to *F* after an extended bottleneck (i.e., mountain goats vs. caribou and white-tailed deer; Figure S2), which further implies that ROHs require both time and demographic change to accumulate (see Gillespie, 2004). Overall, these results support the idea that genome-wide effects of demographic change will not accrue unless given enough time; genome derived *F* and *F_ROH_* are therefore unlikely to align with IUCN conservation status decisions based off census data, similarly noted by Schmidt et al. (2022).

### Mutational Load and Purging

The relationship between *N_e_*, inbreeding, *π*, and mutational load are variable (Grossen et al., 2020; van der Valk et al., 2019; von Seth et al., 2021). Empirical studies on mammals led us to predict that low diversity and high inbreeding do not necessarily result in the accumulation of deleterious alleles (Grossen et al., 2020; Robinson et al., 2018, 2018; van der Valk et al., 2019). Here, we found a much greater number of putatively deleterious mutations in white-tailed deer than in mountain goats and caribou by identifying CE-bound SNPs with GERP. These results were confirmed by the deleterious alleles identified by SIFT that showed a similar gap between white-tailed deer and the other species, regardless of the annotation quality and gene number (Table 1). This means that despite lower diversity and small *N_e_*, caribou and mountain goats have relatively low mutational loads; further, despite a relatively low number of CEs, deer still possessed significantly more CE-bound SNPs indicative of deleterious mutations. Mountain goats, in contrast, were found to have the highest amount of inbreeding, but accumulated the fewest deleterious alleles. This reiterates the finding that inbreeding does not necessarily result in high mutational load; in the case of caribou, if *N_C_* can return (e.g., Peart et al. 2020) or if purging of deleterious alleles can occur (e.g., Huber et al., 2020), the genetic cost to the current decline in *N_C_* could be minimized. This, however, remains an open question and likely population specific, as for example wolves in northern Europe show fixation, not purging, of deleterious alleles (Smeds & Ellegren, 2022).

The efficiency of purging is dependent, in part, on the rate of inbreeding and the genetic load architecture (Hedrick & Garcia-Dorado, 2016;); there is evidence that it may occur more efficiently in the presence of severe bottlenecks (Figure 4b; Grossen et al., 2020). Mountain goats were shown to have low mutational load in conjunction with generally low diversity, while caribou present intermediate profiles. This would be consistent with *N_C_* numbers pre-20^th^ century for caribou, that were presumably much higher (Festa-Bianchet et al., 2011). Reductions in *N_e_* from severe bottlenecks can increase the fitness effects of recessive deleterious alleles by increasing homozygosity; this strengthens the pressure of negative selection and can ultimately lower the mutational load of a population (Garcia-Dorado, 2012, 2015; Glémin, 2003). However, this effect can be insufficient for overcoming cases of extensive homozygosity. Sumatran rhinos for example have a very similar historical *N_e_* and ROH profiles to that of mountain goats (von Seth et al., 2021); while purging might have played some role in some populations in rhinos (von Seth et al., 2021), the loss of habitat combined with overall negative effect of autozygosity and continued presence of high frequency deleterious alleles (e.g., Khan et al., 2021) was likely insurmountable from a population viability standpoint.

Purifying selection should remove SNPs in functionally important regions and constrain these areas against change, which whould generate less SNPs than expected in CEs and lower dN/dS values (Cooper et al., 2005; Davydov et al., 2010; Kryazhimskiy & Plotkin, 2008). The mountain goat’s longer and more numerous CEs, and lowest dN/dS values relative to the other species are suggestive of stronger purifying selection. In contrast, caribou have a smaller *N_e_* than white-tailed deer; thus, purging has either yet to take place or selection is less efficient, or a combination of both. The trajectory of caribou would be predicted to be towards the patterns of mountain goat based on mammalian wide patterns and profiles (Cooper et al., 2005; Huber et al., 2020). It is also important to note that purifying selection reduces nucleotide diversity by eliminating sites under direct selection and those linked through background selection (Charlesworth, 1996; Cvijović et al., 2018), which partially explains patterns of diversity seen here (Figure 1; Buffalo, 2021).

This work has implications for conservation decisions and the expected genomic consequences of numerical population declines. While deleterious alleles may be purged in small, inbred populations, extremely high levels of autozygosity can render this benefit insignificant. Thus, we posit that high mutational load must be accompanied by high ROH to be of conservation concern, similar to what was observed in woolly mammoths (Rogers & Slatkin, 2017). Further, genetic rescue decisions should not rely solely on inbreeding and diversity metrics, especially when their relationships to population viability (Carley et al., 2022) and conservation status (Schmidt et al., 2022) are not straightforward. This is also true for assessing restocking efforts, as we demonstrated that translocations are not necessarily related to population health metrics (Table S2). Collectively, this provides further evidence of the time needed for genome-scale changes to accrue after significant changes in demography. Therefore, we recommend that genomic health metrics like *F_ROH_*, mutational load, and evolutionary constraint be examined alongside classic metrics like *π* and Tajima’ D for a holistic view that can provide more accurate assessments of genome health.

## Acknowledgements

We wish to acknowledge that the work for this study was carried out on the traditional territory of the Mississauga (Michi Saagiig) Anishinaabeg people. This work was supported by a Natural Sciences and Engineering Research Council Discovery Grant, Canada Foundation for Innovation: John R. Evans Leaders Fund, and Compute Canada Awards to ABAS; EW was supported by three NSERC USRAs. We thank Daria Martchenko, Morgan Dedato, and Camille Kessler for help with data acquisition. Sample providers are listed in Dedato et al. (2022), Martchenko & Shafer (2022), and Kessler et al. (2022). We thank three anonymous reviewers for comments that strengthened this manuscript.

## Data Availability and Benefit Sharing

All genome assemblies can be found on GenBank (details in Appendix Table S1). Re-sequencing data has been submitted to SRA and accession numbers are pending. Bash and R scripts for all analyses are available on GitLab: https://gitlab.com/WiDGeT_TrentU/undergrad-theses/-/tree/master/Wootton_2022

Benefits Generated: This research addresses important questions about the genomic health of populations of priority concern and illustrates relationships between genomic health and long-term demographic history. Collaborations were formed with researchers across North America, and all major collaborators are included as co-authors. All contributors are listed in the Acknowledgements. We are committed to institutional capacity-building and have shared the results of our research with our collaborators and the broader scientific community.

## Author Contributions

ABAS and EW designed the research project. EW conducted all analyses. ABAS and EW wrote the manuscript using comments and contributions from all coauthors.

## Supplemental Information

**Table S1:** GenBank assembly accession numbers for each species used by GERP.

| Species | GenBank Assembly Accession |
| --- | --- |
| *Equus caballus* | GCA_002863925.1 |
| *Equus asinus* | GCA_016077325.2 |
| *Tapirus terrestris* | GCA_004025025.1 |
| *Diceros bicornis* | GCA_004027315.2 |
| *Ceratotherium simum* | GCA_000283155.1 |
| *Camelus dromedarius* | GCA_000803125.3 |
| *Camelus bactrianus* | GCA_000767855.1 |
| *Hippopotamus amphibius* | GCA_004027065.2 |
| *Tragulus javanicus* | GCA_004024965.2 |
| *Moschus moschiferus* | GCA_004024705.2 |
| *Moschus chryogaster* | GCA_006461725.1 |
| *Bos taurus* | GCA_002263795.2 |
| *Connochaetes taurinus* | GCA_006408615.1 |
| *Ovis canadensis* | GCA_004026945.1 |
| *Giraffa camelopardalis* | GCA_017591445.1 |
| *Antilocapra americana* | GCA_007570785.1 |
| *Alces alces* | GCA_007570765.1 |
| *Cervus elaphus* | GCA_910594005.1 |
| *Phacochoerus africanus* | GCA_016906955.1 |
| *Sus scrofa* | GCA_000003025.6 |
| *Rangifer tarandus* | GCA_019903745.1 |
| *Oreamnos americanus* | GCA_009758055.1 |
| *Odocoileus virginianus* | GCA_014726795.1 |

**Table S2:** Correlation matrix between genomic health metrics and translocation data for white-tailed deer.

| | $F_{ROH}$ | $F$ | Fraction of deleterious CEs | Fraction of deleterious genes |
| --- | --- | --- | --- | --- |
| **Stock start** | 0.032 | 0.127 | 0.076 | -0.225 |
| **Stock end** | 0.372 | 0.392 | 0.205 | 0.347 |
| **Population low** | -0.182 | -0.059 | -0.177 | 0.308 |
| **Number stocked** | -0.102 | -0.062 | 0.162 | -0.103 |
| **Number of states** | -0.161 | -0.164 | -0.093 | 0.096 |

**Figure S1:**
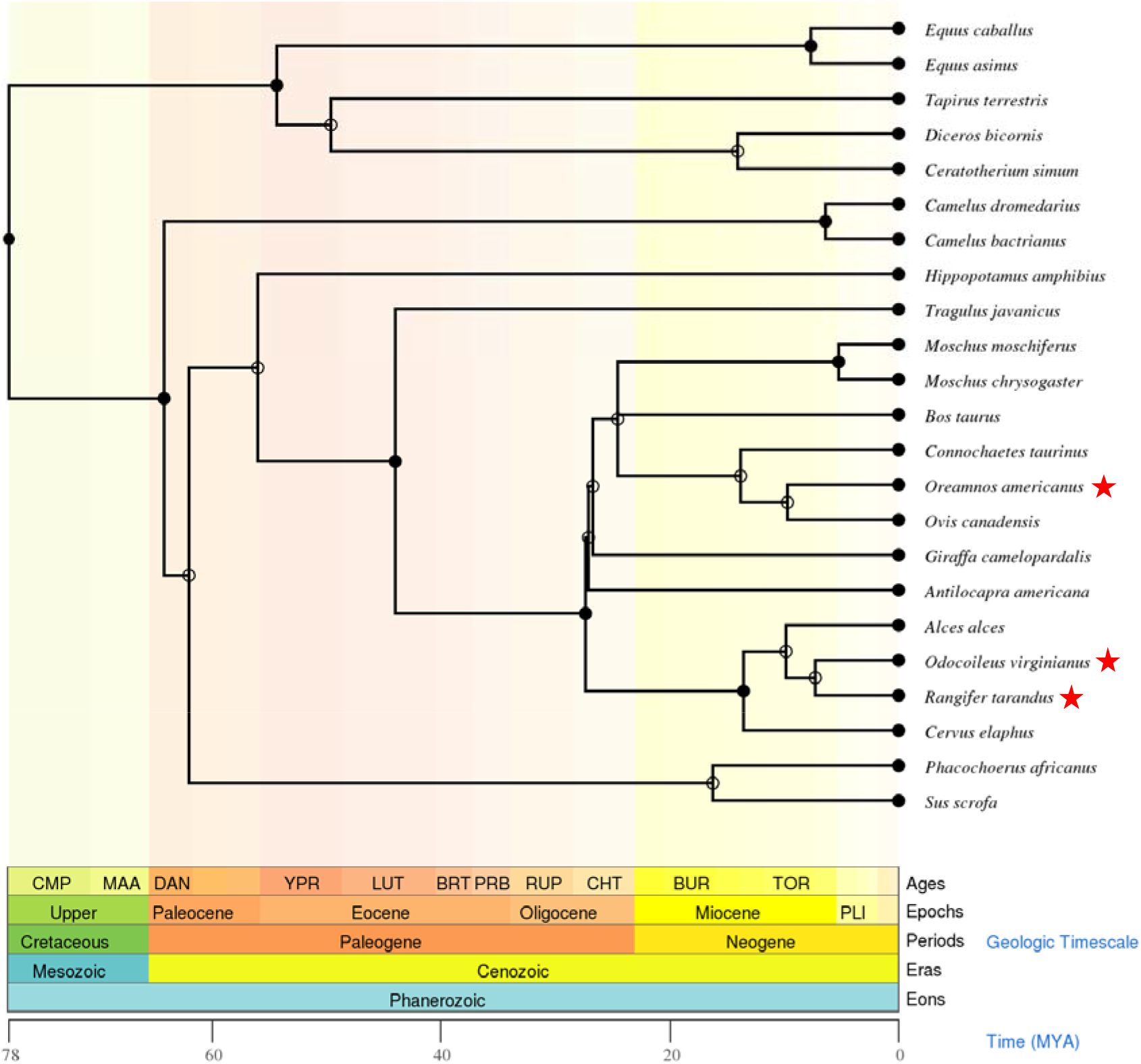
Phylogenetic tree generated by TimeTree showing all 23 ungulate species analysed by GERP++ and estimated times of divergence in MYA (Kumar et al., 2017). Study species with sample data are starred in red (*Odocoileus virginianus, Rangifer tarandus*, and *Oreamnos americanus*).

**Figure S2:**
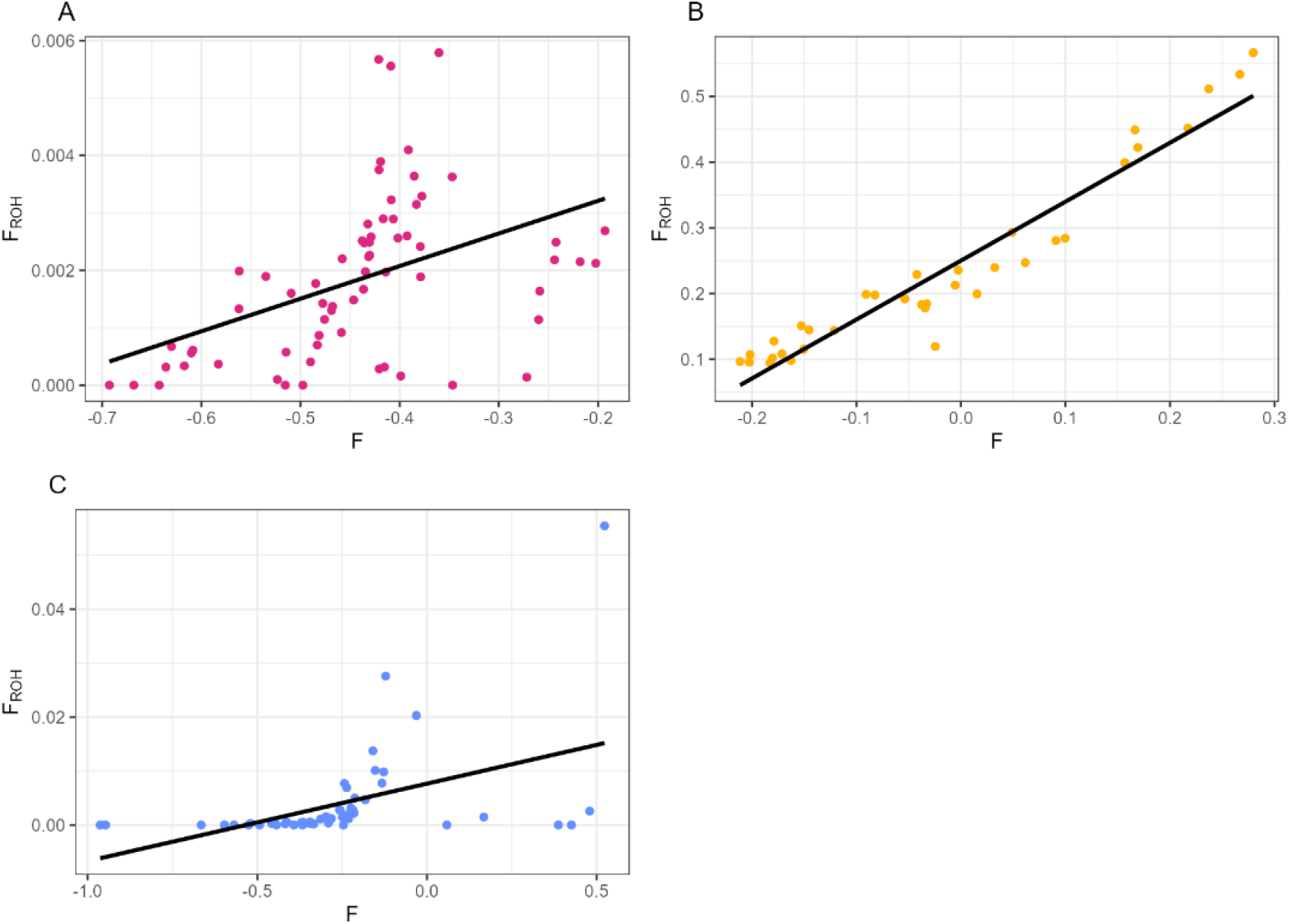
(A) Scatterplot and linear regression line showing the correlation between *F* and *F_ROH_* in caribou (Pearson’s *r* = 0.199, *p* < 0.05). (B) Scatterplot and linear regression line showing the correlation between *F* and *F_ROH_* in mountain goats (Pearson’s *r* = 0.918, *p* < 0.05). (C) Scatterplot and linear regression line showing the correlation between *F* and *F_ROH_* in white-tailed deer (Pearson’s *r* = 0.213, *p* < 0.05).

**Figure S3:**
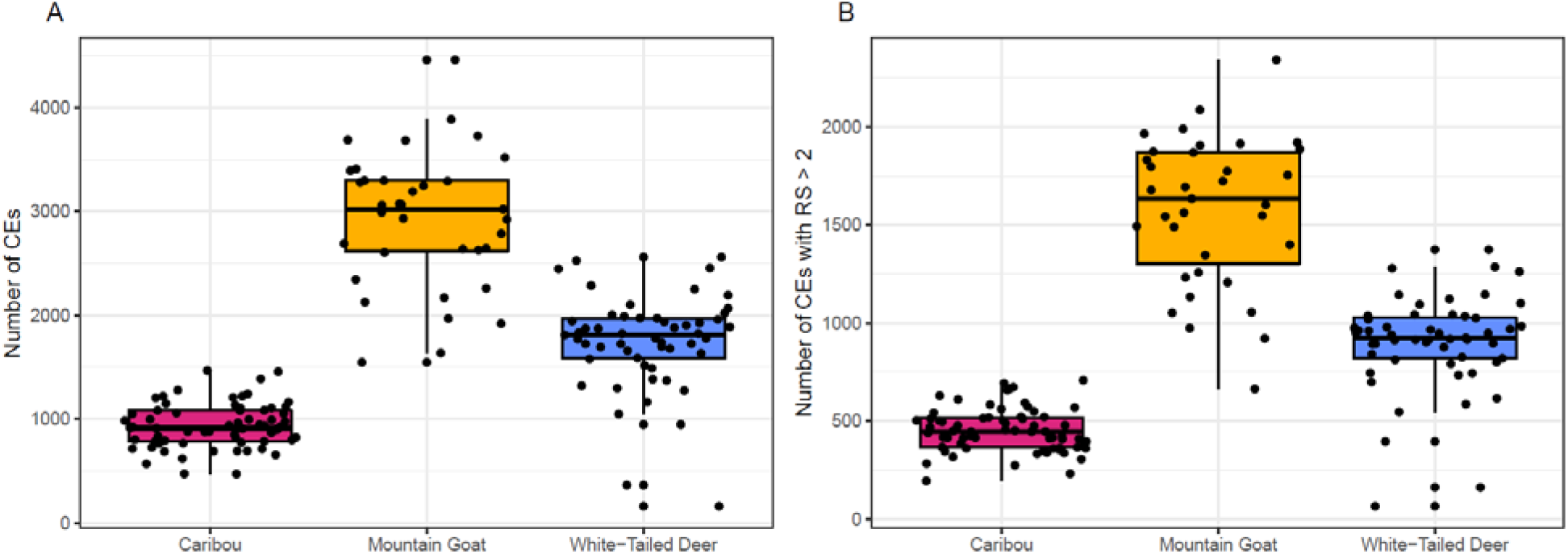
(A) Boxplot showing the number of constrained elements (CEs) identified per species by GERP++’s gerpelem function. (B) Boxplot showing the number of highly constrained CEs (RS >2) identified in each species by gerpelem.

**Figure S4:**
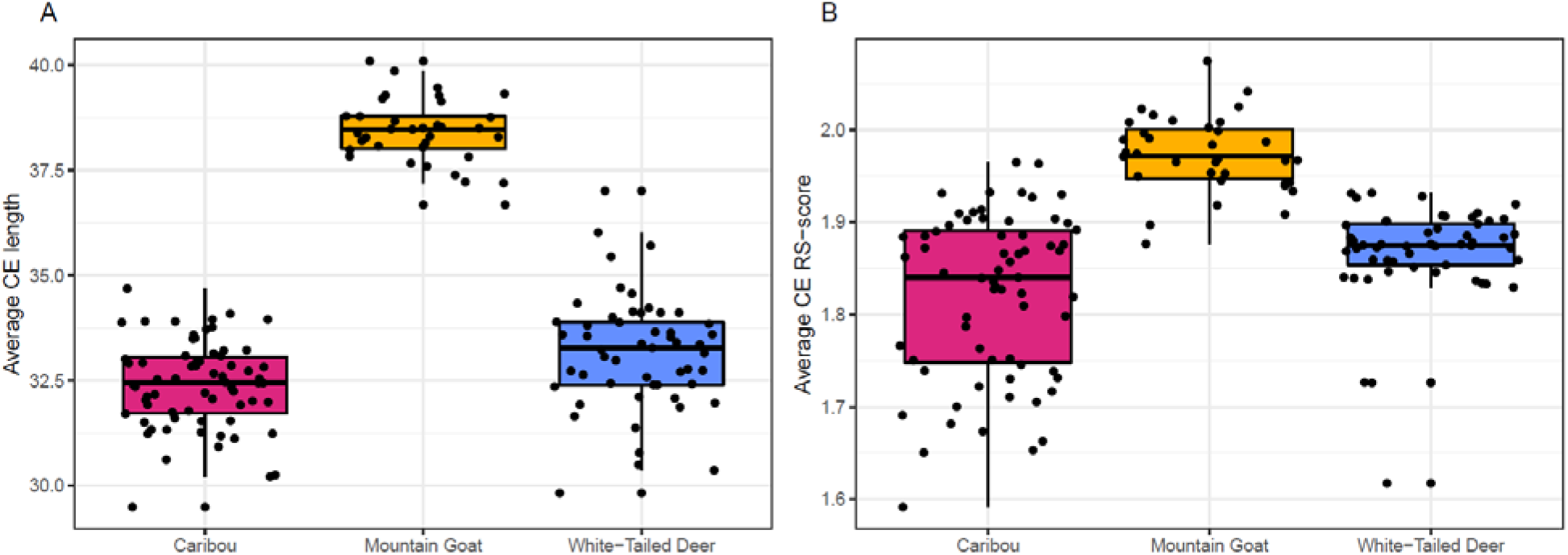
(A) Boxplot showing the average length of constrained elements (CEs) identified per species. (B) Boxplot showing the average intensity of constraint (RS-score) of CEs identified in each species.

**Figure S5:**
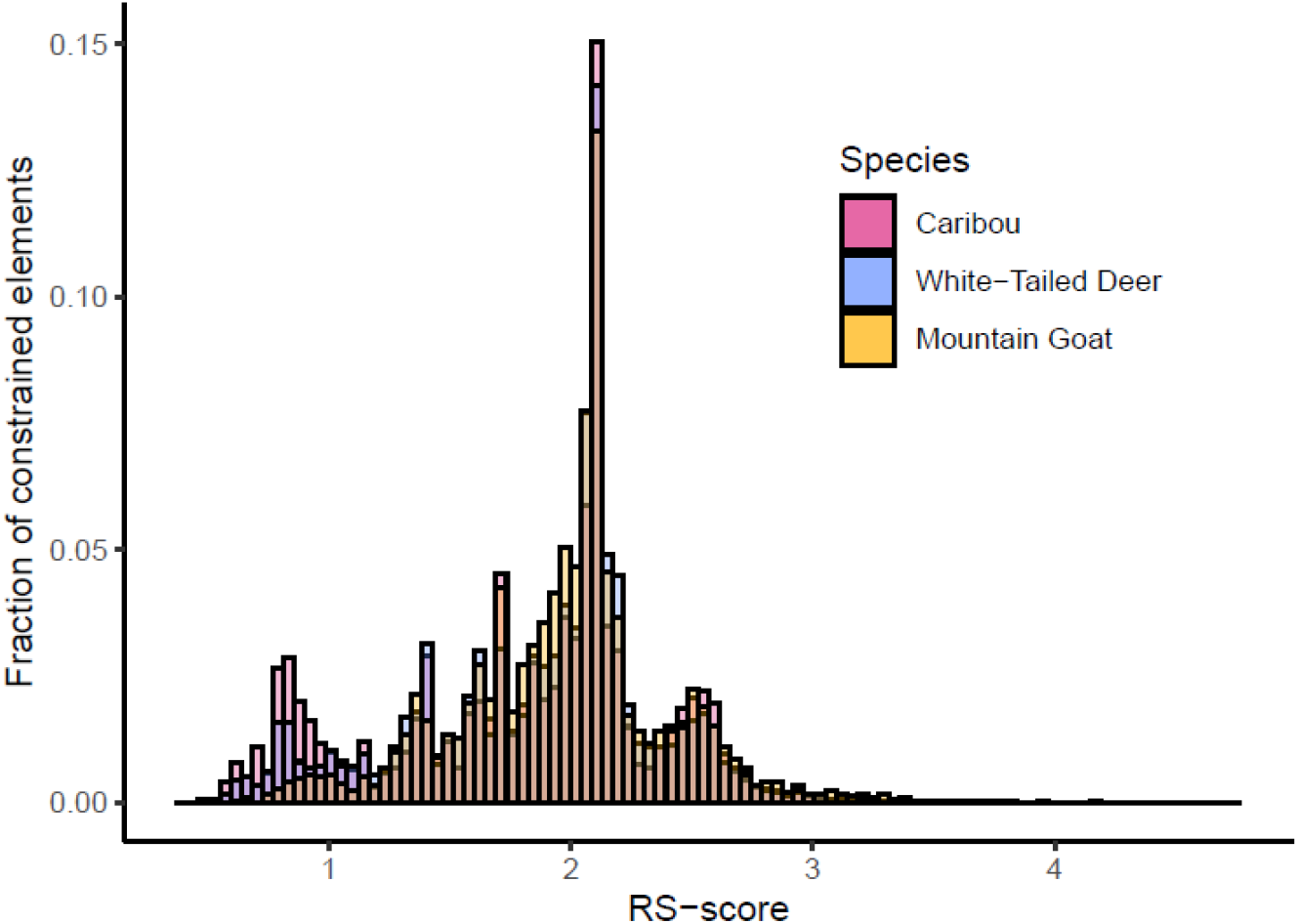
Histogram showing the fractional distribution of constrained element (CE) RS-scores.

**Figure S6:**
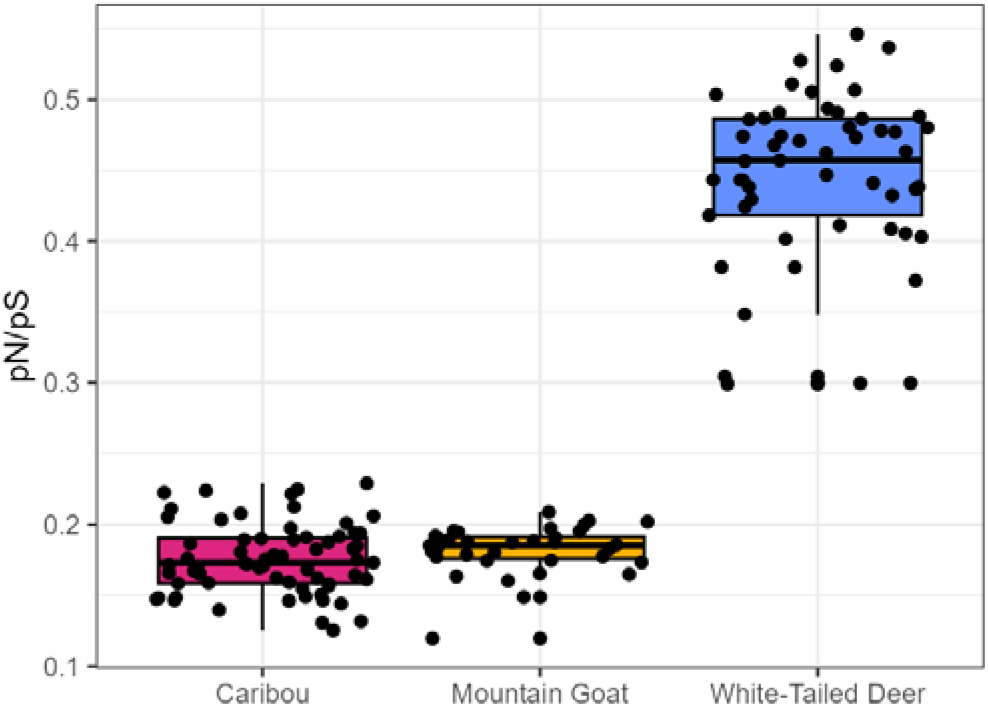
Boxplot showing the ratio of nonsynonymous to synonymous mutations (pN/pS) identified by SIFT in each species.

**Figure S7:**
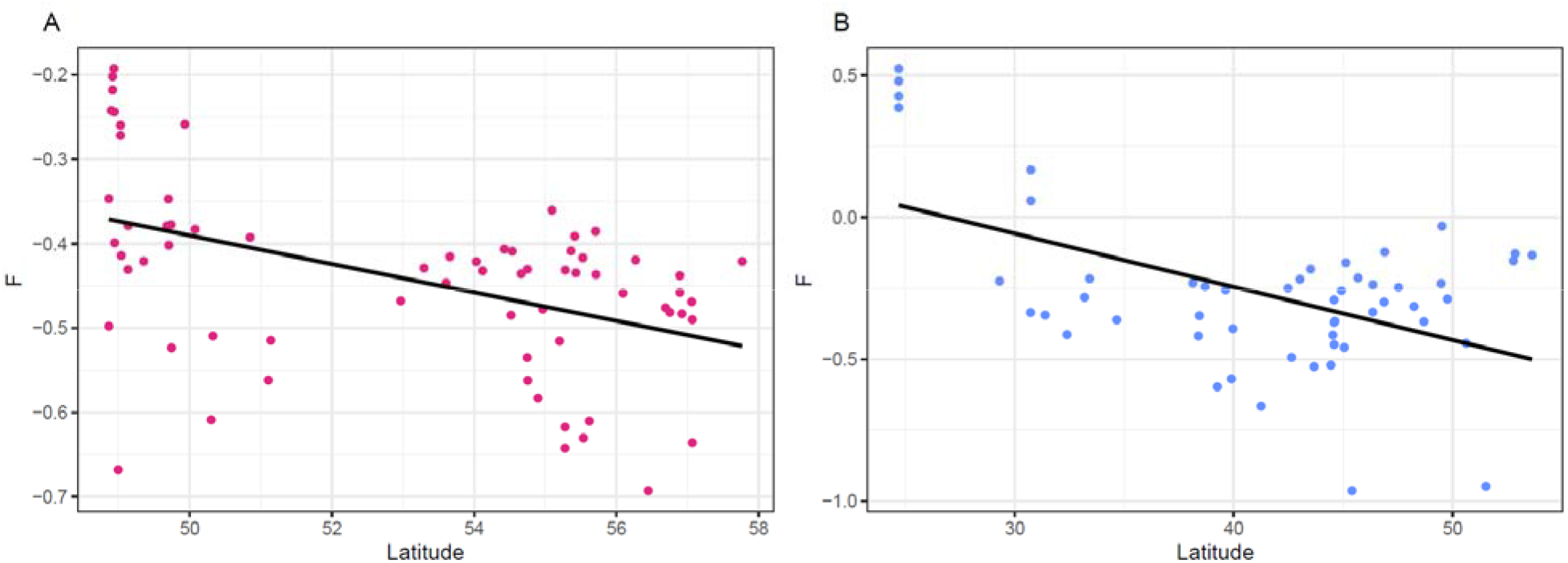
(A) Scatterplot and linear regression line showing the correlation between *F* and latitude in caribou (Pearson’s *r* = −0.466, *p* < 0.05). (B) Scatterplot and linear regression line showing the correlation between *F* and latitude in white-tailed deer (Pearson’s *r* = −0.513, *p* < 0.05).

